# Exploring ancestry inference of the Middle East

**DOI:** 10.1101/2024.08.15.607793

**Authors:** Noah Herrick, Mirna Ghemrawi, Sylvia Singh, Rami Mahfouz, Susan Walsh

## Abstract

The capability to infer biogeographic ancestry with curated panels of ancestry informative markers (AIMs) is a critical component to DNA intelligence. There are many AIM panels that resolve population differentiation at a continental level. Of late, several studies have directed marker discovery to the Middle East because of the difficulties for AIM panels to resolve this region amongst populations in Eurasia. The AIM discovery process has remained largely unchanged, except for the most recent additions of whole-genome sequence (WGS) data repositories which now include Middle Eastern individuals. Here, the latest WGS data from 1000 Genomes Project and Human Genome Diversity Project was paired with novel Middle Eastern population data from Lebanon for AIMs discovery. An unbiased genetic clustering approach was employed for selecting population clusters for allelic frequency comparisons. Two candidate AIMs were reported, compared, and evaluated together with the autosomal AIMs from the VISAGE Enhanced Tool. These comparisons involved a validation dataset from Middle Eastern WGS data published by the Wellcome Sanger Institute and resulted in slight gains of Middle Eastern ancestry proportions for several Middle Eastern samples with varying levels of co-ancestries. The validation samples also underwent an unsupervised worldwide ADMIXTURE analysis alongside previously mentioned WGS datasets using nearly two million markers (r2 < 0.1) to establish a ‘ground truth’ population membership. Lastly, a novel application of the deep learning dimensional reduction algorithm ‘popVAE’ is provided as an open-source web tool to illustrate the AIM panels variance among these population clusters within two dimensions for easy global ancestry visualization in addition to providing a closest population membership metric.

## 1. Introduction

Intelligence-driven methodologies have seen an increase in use over the last decade, spanning the prediction of physical appearance [1-6], the inference of biogeographical ancestry [7-13], and the investigative genetic genealogy search of familial relationships [14-16]. As research into the choice of markers is explored and panels continue to be developed, there is one challenge that is typically encountered in a forensic setting: DNA. Specifically, the limitation in quantity and quality of DNA obtained from the low amounts of biological material present [17]. In an ideal scenario, whole-genome sequencing or genome-wide variant panels are chosen to generate data. However, DNA limiting factors force researchers to fine-tune smaller variant panels and their designs to yield the most genotypic information from as little DNA as possible, all while achieving their intelligence information goal and accuracy standard.

Genetic panels containing small sets of ancestry informative markers (AIMs) with high population divergence are typically used for accurate biogeographical ancestry (BGA) inference. These can range from combinations of bi-allelic to multi-allelic autosomal markers, microhaplotypes, and lineage markers using Y-chromosomal and mitochondrial typing. The VISAGE Enhanced Tool [18] (VISAGE-ET-AA), a combined panel for appearance prediction and ancestry inference, stands out among many forensic ancestry panels designed in the last decade [19-28] as the newest and most exhaustive for continental BGA inference. It is made up of 184 appearance markers and 226 ancestry markers which are subset into 104 autosomal SNPs, 85 Y-SNPs, 16 X-SNPs, and 21 microhaplotypes. VISAGE-ET-AA is an extension of the VISAGE Basic Tool [26] (VISAGE-BT-AA), which consisted of 115 AIMs and all 41 markers from the HIrisPlex-S panel [6]. Notable aspects of VISAGE-ET-AA include the considerable effort dedicated to expanding the number of predicted phenotypic traits, as well as the incorporation of an increased number of Middle Eastern (ME) informative variants intended to enhance the resolution of BGA inference for the ME region.

A geographically central region between the continents of Europe, Central/South Asia, and Africa, the Middle East is of particular interest due to influences from agriculture and the neolithic revolution [29], population migrations [29, 30], and cultural and religious influences [31, 32]. The marginal genetic divergence between the Middle Eastern and Eurasian population groups of Europe, North Africa, and South Asia exists, in part, from the heavy influence of gene flow throughout history [33] and has proved challenging [22, 34-36] to accurately make inferences between these genetically similar groups. Sequencing data for the Middle East from HGDP-CEPH [37] and the Sanger Wellcome Institute [38] has since provided an opportunity for improved AIM discovery and more accurate BGA inference [18] of the region. For example, it has led to an increase in ME-specific markers in the VISAGE-ET-AA autosomal variant set, which now contains 28 ME-specific variants out of the 104 total. Additionally, over half (63) of the 104 autosomal BGA variants representing seven global populations were novel and unique to the VISAGE-ET-AA panel. This recapitulates the strength of marker discovery with sequencing and novel population data in that work [18]. Exploration of other unique populations in the region, such as Lebanon, may further assist in increasing the list of ME-informative markers that could further improve resolution of this region and complement current autosomal panels being used.

Lebanon occupies a central location within the Middle East, withstanding as the crossroads of the Arabian Peninsula and the Mediterranean region. In addition to its notable geographical location, an exploration into the population genetics of Lebanon [39] has revealed a genetic continuity from a Canaanite history. It was observed that modern-day Lebanese individuals may share up to 93% of their ancestry with this ancient civilization which is sometimes also referred to as Phoenicia. These were indigenous populations thousands of years ago who settled in the Near East [39], made-up of present-day Jordan, Syria, Palestine, and Lebanon. Evidence of strong genetic continuity with ancestral populations of the Middle East further supports including the Lebanese subpopulation in ME AIM discovery studies.

Classical approaches to search and evaluate BGA autosomal variants such as informativeness for assignment metric (I_n_) [8], allele-frequency differential between populations (δ) [7], or fixation index (FST) [40, 41] are typically used to rank AIM candidates for their utility in ancestry resolution and inference. Each of these calculations are based on allele frequencies at their root, and studies attempting to identify AIM candidates for a particular geographical region benefit from the addition of novel population datasets to help solidify allelic frequency differences. However, it is important to note that these results can be highly influenced by the definition of the populations [12]. This constitutes either a geographical supervised approach or a genetic unsupervised approach. The former alludes to categorizing individuals based on known, self-reported ancestry. The latter is based on popular model-based clustering algorithms and programs such as ADMIXTURE [42] and STRUCTURE [43], which can be used to define the cluster or region (and the individuals that make up that cluster/region) in order to rank the candidate AIMs for optimal population separation in ancestry inference.

There are several ways to assess the performance of BGA inference using small sets of AIMs. The first step is establishing the most probable ancestry or ‘ground truth’ of the reference set. This can be established either i) using a geographical designation or collection site label and/or ii) a thorough exploration of their population structure using whole-genome and genome-wide variant information in an unsupervised genetic approach with reference data [44]. Once the ground truth is known, the second step is to assess the BGA inference accuracy with robust tools. One of these tools is a naïve Bayesian classifier, Snipper [10, 45], that utilizes a likelihood ratio analysis to provide the most likely ancestry in relation to a reference training set to assess classification accuracy. Another tool that is widely utilized for testing AIMs is the model-based genetic clustering program, STRUCTURE, which can demonstrate an AIM panel’s ability to differentiate ancestry across broad (or best K) continental clusters/regions. Principal components analyses (PCA) have also been used in population genetic studies [46] to visualize where the unknown individuals cluster in a dimensionally reduced genetic ancestry space [11, 47-52].

In this study, we supplemented the marker discovery process for Middle Eastern AIMs with a novel population dataset (n=190) from Lebanon in combination with 1000 Genomes Project (1KG) and Human Genome Diversity Project (HGDP) data, using unsupervised genetic clustering to define the population groups. We then assessed the classification performance of the VISAGE-ET-AA autosomal AIM panel for Middle Eastern ancestry inference with and without the inclusion of our novel ME AIMs using Snipper, STRUCTURE and PCA on a Middle Eastern population test set from the Sanger Wellcome Institute [38]. Finally, in an effort to improve ancestry inference visualization overall using Forensic AIM panels, we implemented and adapted a deep learning, non-linear variational autoencoder (VAE) algorithm called popVAE [53] to highlight the visual improvements seen for ancestry inference with AIMs when moving to a deep learning derived 2D latent space capable of capturing global and local genetic variance.

## 2. Material and Methods

### 2.1. Genetic Datasets

#### 2.1.1. Lebanese population data

220 samples were collected from study volunteers in Beirut, Lebanon after Institutional Review Board (IRB) approval from the American University of Beirut (PALM.RM.27). After the removal of related individuals and several participants with questionnaire data stating non-Lebanese ancestry, a final number of 190 unrelated individuals with reported Lebanese ancestry were used in this study. Analysis of this data was conducted under additional IRB approved protocol IU-IRB #1409306349 within the United States. Written informed consent documents were obtained from each of the donors including questionnaire data stating information on the place of birth, including both paternal and maternal sides, of each participant.

DNA was extracted from the saliva samples collected using an in-house salting out method and then quantified on an Invitrogen Qubit Fluorometer in combination with a DNA High Sensitivity Assay Kit (Fisher Scientific International Inc. Hampton, NH, USA). DNA was genotyped using the Illumina (Illumina, Inc. San Diego, CA, USA) Infinium Multi-Ethnic Global-8 v1 array (MEGA) consisting of 1,733,365 genome-wide markers. Genotyping was performed by the University of Chicago’s DNA Sequencing & Genotyping Facility (Chicago, IL). The raw microarray BeadArray data was obtained from IDAT files corresponding to each sample. These files were transformed into one variant call format (VCF) file for all 190 samples using the genomic data processing tool, *Iliad* [54]. The *Iliad* SNP array workflow also performed several variant cleaning steps: converting Illumina loci names for conventional reference SNP cluster identifiers (rsIDs), filtering for dbSNP annotation file rsID overlap, and keeping variants above GenTrain and ClusterSep array quality thresholds of 0.7 and 0.45, respectively. The final optimized VCF contained 1,686,450 SNPs.

#### 2.1.2. Reference and test data

Population genetic data was obtained from the 1000 Genomes Project [55] (1KG), the Human Genome Diversity Project [37] (HGDP), and the Wellcome Sanger Institute [38] (SangerME) in the form of WGS data representing seven super populations: Africa (AFR), America (AMR), East Asia (EAS), Europe (EUR), Middle East (ME) Oceania (OCE), and South Asia (SAS). Both 1KG and HGDP reference datasets were obtained in CRAM format and processed using the stored sequence module of *Iliad* [54]. The SangerME dataset was retrieved from the NCBI Sequence Read Archive (SRA) using *Iliad*’s raw sequence module configured for SRA toolkit [56] (v3.0.2) data retrieval and was reserved for testing purposes only.

### 2.2. Middle East (ME) AIM discovery

#### 2.2.1. Genetic data quality control & cluster assignment

The Lebanese and reference 1KG (30X) and HGDP (30X) data were combined which resulted in a final VCF of 3,522 samples and 1,307,875 SNPs. The combined dataset was filtered to remove variants with less than 5% missingness using BCFtools [57] (v1.17) and then converted to PLINK2 [58] (v2.00a4LM) where additional quality control (QC) measures were applied: LD pruning (r2<0.1), HWE, and a 1% MAF threshold. This reduced the number of variants to 194,321 SNPs. Principal components analysis (PCA) was performed using PLINK [59]. The variance of our dataset was visualized using ggplot2 [60] within the R (v3.5.1) environment. The package Ellipses [60], based on the Khachiyan ellipsoid algorithm, was used to depict labelled geographical population data with calculated cluster boundaries and to provide a PC 1-2 averaged coordinate (square) within each ellipse (Figure 1A).

**Figure 1.**
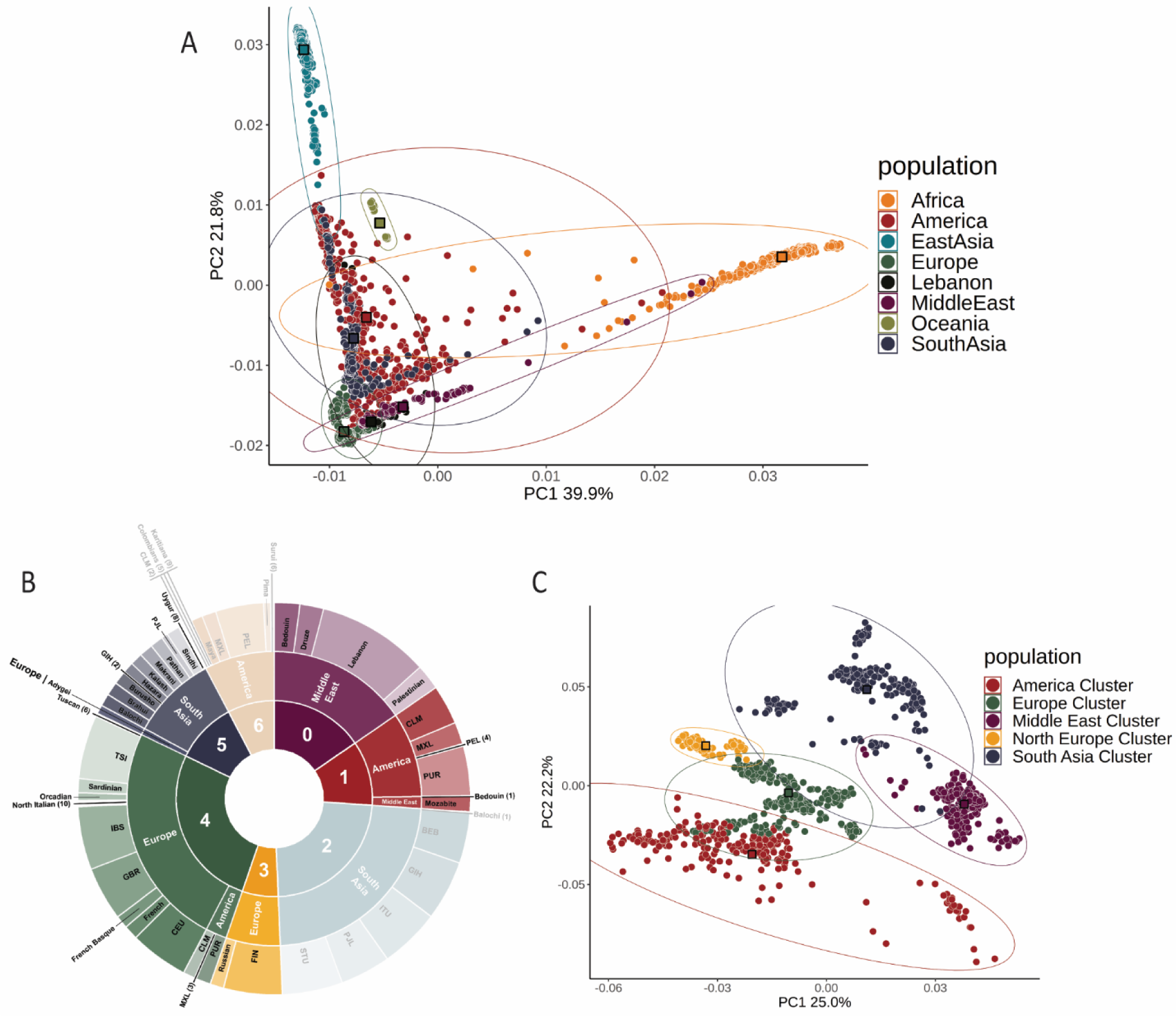
Optimization of ME-AIM discovery dataset. A) PCA plot of population data from 1KG (n=3332), HGDP (n=828), and Lebanon (n=190) on markers in linkage equilibrium at an r2 threshold of less than 0.1 (n=194,321). Colors are assigned by their geographical label. B) Distribution of population data after K-means cluster assignment using ADMIXTURE ancestry components (East Asia, Africa, and Oceania were removed prior to this analysis). Colors represent the cluster assigned by K-means. The faded sections of the plot represent clusters (2 and 6) that were removed prior to downstream marker discovery steps. C) PCA plot of remaining clusters (0: Middle East, 1: America, 3: North Europe, 4: Europe, 5: South Asia) that were used in pairwise comparisons for marker discovery (SNP=754,826). Colors are assigned by their K-means cluster label.

ADMIXTURE [42] (v1.3) was used for maximum likelihood estimations in an unsupervised manner to assign ancestral component percentages to each of the individuals selected for targeted ME AIM discovery. The input format required by ADMIXTURE was a PLINK (v1.90b7) binary (.bed) fileset. The data was previously pruned of SNPs in linkage disequilibrium (LD) at an r^2^ threshold under 0.1 and met the linkage equilibrium assumption of ADMIXTURE. Models of K:2-16 were applied to the combined dataset where ‘K’ was the number of possible populations for ADMIXTURE to assign each of the 2,051 individuals. Each of the 15 models were replicated 10 times to find the parameter standard errors declared by the ‘-B10’ flag for bootstrapping procedure. Ten-fold cross-validation was applied to each model using the ‘--cv=10’ flag. The model chosen to represent the most optimal number of population clusters displayed the lowest cross-validation (CV) error. K-means clustering algorithm from the sklearn.cluster package in Python (v3.7.12) was then applied to the most optimal model K. The ancestral component percentages estimated by ADMIXTURE were used as input to classify individuals into an appropriate cluster of most genetic similarity. This method relied on an unsupervised ancestral component clustering approach rather than geographical population labels. The CLUMPAK [61] server was used for visualization of the best K ancestral component proportions.

Genetic differentiation tests were performed to reduce the number of genetic clusters for downstream analyses to those most geographically and genetically proximal to the Middle East. These metrics were determined by minimal mean FST values and further supplemented with the minimal variance observed in PCA plots. The mean FST values among the clusters were calculated using the Weir and Cockerham F-statistic [41] in PLINK2 ‘--fst method=wc’.

#### 2.2.2. Genetic cluster allelic comparison

The absolute value of the allele frequency difference (delta; δ) per SNP was calculated between clusters. These clusters were renamed based on the single region with the highest ancestry proportion observed in each cluster (i.e., Middle East, America, Northern Europe, Europe, and South Asia clusters). Allele frequencies of all markers that passed QC (n = 754,826) were reported with BCFtools plug-in ‘+fill-tags’ and δ was calculated using in-house python scripts. Variants were examined using dbSNP variation viewer and those positioned in problematic regions of alignment (e.g. if the SNP was surrounded by annotated insertions and deletions) were removed from the candidate list.

### 2.3 ME Ancestry inference using known AIM marker panels

#### 2.3.1. Ancestry cluster assignment for training and test data

An in-depth, unsupervised ADMIXTURE analysis was performed on the full set of n=137 test individuals from the SangerME dataset [38] that included reference samples from 1KG (30X; n=2,504 unrelated individuals), and HGDP (30X; n=828). All whole-genome sequencing (WGS) samples were processed with the *Iliad* [54] suite of Snakemake [62] workflows which calls 120,046,375 variants using the NYGC annotation files [63]. These variants were filtered for 5% missingness, LD (r2<0.1), HWE, and a 1% MAF threshold leaving 1,712,516 autosomal variants for unsupervised ancestry assessment using ADMIXTURE [42]. Models of K:6-10 with 10-fold cross-validation were applied to find the best K with the lowest CV error. To illustrate how samples flow from one K into another, the package “ggalluvial” [64] was used in the R environment [65] to generate a sankey diagram using a “ggplot2” [66] extension. The reference populations were filtered for “Anchor” populations which consisted of samples identified from 1KG and HGDP that presented proportions at or above the average main ancestry component of their cluster at the best K. The Anchor reference clusters were kept for BGA inference training.

#### 2.3.2. Snipper and STRUCTURE analysis

The panel of autosomal AIMs from the VISAGE-ET-AA [18] were used for BGA inference. The candidate variants from this study were also included for comparison. Although this VISAGE-ET-AA panel originally consisted of 104 autosomal variants, only 102 AIMs were used in this study due to noted issues [13, 18] with the variants rs3857620 and rs17287498. Genotypic information was obtained for the VISAGE-ET-AA variants from the WGS datafiles as described above. Anchor reference clusters were used as training data for Snipper [10, 45] (v3), the naïve Bayesian classifier, to assess the ancestry prediction outcomes of the SangerME independent test samples. Scripts to run the classifier were downloaded from the snipper website found at (http://mathgene.usc.es/snipper/offline_snipper.html), and included the use of packages ““klaR” [67], “openxlsx” [68], “caret” [69], “ggplot2” [66] in the R [65] environment. A supervised STRUCTURE [43] analysis was also performed using the Anchor reference clusters (POPFLAG = 1) on the 137 SangerME test set (POPFLAG = 0) at 100,000 burn-in steps and 100,000 MCMC steps using the VISAGE-ET-AA AIM panel (n=102). A second STRUCTURE run was performed with the same parameters but included the additional candidate ME-AIMs from this study. The ancestry components of the SangerME samples were plotted using CLUMPAK [61].

### 2.4. Ancestry inference visualization

PCA and ‘popVAE’ [53], a deep learning variational autoencoder (VAE), were two independent methods used to visualize and compare the genetic variance of the AIM panels tested. The popVAE program was downloaded from the Github repository found at (https://github.com/kr-colab/popvae) and installed in the form of a conda environment [53]. Both biallelic and multiallelic genotypes were normalized between 0 and 1 prior to input. The missing genotypes for samples were handled by inputting the average genotype across each population cluster group. To optimize model building, the training and testing sets were selected solely from the 1KG and HGDP Anchor data (Supp. Table 1). They were balanced and adjusted for a random selection of 5 samples tested from each population cluster. A total of 20 models were generated by setting seed flags of 1-20. After examination of the scatterplots and learning curves, the encoder model parameters were saved for the best model that showed a balance between validation and training loss, thus reducing overfitting. To assess the model’s performance on an independent test set, AIM genotypes for each individual within the SangerME dataset were loaded with an added “unknownsfile” flag and their latent space distribution within the 1KG and HGDP Anchor ancestry space was predicted. Models were generated and samples plotted via projection for the VISAGE ET (n=102) panel together with the ME-AIMs best candidates from this study. A 2D Euclidean distance metric was generated by calculating the difference between the unknown individuals predicted latent co-ordinates and the centroid point of each population cluster to indicate ancestry inference. Although the predictions can handle missing genotype data, specifically in the form of a blank value or NA by substituting the average allele frequency of the Anchor dataset as the missing variant, it is encouraged to keep the amount of missing data to a minimum for the best possible results.

**Table 1.** Snipper likelihood ratio table for incorrect classification comparisons between ET and ET+2.

| Sample | Deep<br>ADMIXTURE<br>Maximum K | AIMs<br>Panel | Predicted<br>Population | Middle<br>East<br>Cluster | South<br>Asia<br>Cluster | Europe<br>Cluster | Africa<br>Cluster |
| --- | --- | --- | --- | --- | --- | --- | --- |
| APPG7555904 | Africa<br>Cluster | ET | Middle East Cluster | 0.99 | 0 | 0 | 0.01 |
|  |  | ET+2 | Middle East Cluster | 0.99 | 0 | 0 | 0.01 |
| APPG7836404 | Middle East<br>Cluster | ET | Africa Cluster | 0 | 0 | 0 | 1 |
|  |  | ET+2 | Africa Cluster | 0 | 0 | 0 | 1 |
| APPG7555903 | Middle East<br>Cluster | ET | South Asia Cluster | 0.47 | 0.53 | 0 | 0 |
|  |  | ET+2 | Middle East Cluster | 0.87 | 0.13 | 0 | 0 |
| APPG7555888 | Middle East<br>Cluster | ET | Europe Cluster | 0.36 | 0 | 0.64 | 0 |
|  |  | ET+2 | Europe Cluster | 0.03 | 0 | 0.97 | 0 |
| APPG7555914 | Middle East<br>Cluster | ET | Middle East Cluster | 0.86 | 0.14 | 0 | 0 |
|  |  | ET+2 | South Asia Cluster | 0.27 | 0.73 | 0 | 0 |
| APPG7555885 | Middle East<br>Cluster | ET | South Asia Cluster | 0 | 1 | 0 | 0 |
|  |  | ET+2 | South Asia Cluster | 0 | 1 | 0 | 0 |
| APPG7555872 | Middle East<br>Cluster | ET | South Asia Cluster | 0 | 1 | 0 | 0 |
|  |  | ET+2 | South Asia Cluster | 0 | 1 | 0 | 0 |

In addition, we developed a user-friendly web tool that integrates these modifications with the popVAE [53] algorithm to more easily provide this trained AIM model to the community. The visualization utility of this tool was also assessed with global population samples from Ruiz-Ramírez et al. (2023)^18^. Samples with a greater than 90% genotype coverage (n=368) were included in this assessment. More information on this tool can be found at https://walshlab.indianapolis.iu.edu/pages/tools.html.

## 3. Results

### 3.1. Middle East candidate AIMs discovery

Participants from the Middle Eastern subpopulation, Lebanon (n=190), played a pivotal role in supplementing currently available Middle Eastern data in order to further explore the region’s degree of genetic variation from nearby regions. The additional data improved the Middle Eastern sample size from HGDP by over two-fold for discovery amongst 1,307,875 variants that were genotyped within the Lebanese cohort. Upon initial exploration of the population data when combined with 1KG and HGDP (total n=3,522), PCA and the Khachiyan ellipsoid algorithm (Figure 1A) illustrated that individuals from Africa, East Asia, and Oceania could be removed from further analyses as they were the most genetically distinct from our Middle East regional data. It is also known that these continental populations are more geographically distant than the populations retained for further analyses. There were 2,051 individuals from the remaining population groups for marker discovery which included the Middle East, America, Europe, and South Asia. Although pre-defined population labels have been reported to be substantially informative for BGA genetic groupings [70], we performed an unsupervised, genetic-based population clustering approach on all of the remaining individuals to mitigate genetic noise that can arise from using only geographic labels. The results from ADMIXTURE [42] displayed the lowest cross-validation error at K=7 (Supp. Fig. 1). The K-means algorithm clustered the Lebanese population with other Middle East individuals. Visualization of the ancestral component proportions of K=7 can be seen in Supplemental Figure 2. A full breakdown of the subpopulations, as they clustered, can be found in Figure 1B. Finally, a PCA plot (Figure 1C) highlights the regional centroids of genetic clusters that were used to identify candidate ME-AIMs. Cluster 2 (made up of subpopulations from South Asia) and Cluster 5 (made up of subpopulations from America) were removed from downstream analyses as they did not overlap with the Middle East ellipsoid and showed the largest Weir and Cockerham F-statistic [41] (Supp. Fig 3), indicating they were genetically distinct from the Middle East cluster and therefore may not assist in finding markers to help resolve the Middle East from closer surrounding regions.

ME-AIM discovery consisted of four cluster comparisons: Middle Eastern Cluster (0) vs. American Cluster (1), Middle Eastern Cluster (0) vs. Northern European Cluster (3), Middle Eastern Cluster (0) vs. European Cluster (4), and Middle Eastern Cluster (1) vs. South Asian Cluster (5). Delta (δ) scores were calculated for the four cluster comparisons along 754,826 genotyped variants available in the ME-AIM discovery dataset. The top 500 markers from each pairwise comparison (δ score > 0.3) were explored and quality checked. Some of these markers provided ME differentiation for several overlapping comparisons (Supp. Fig. 4). Two variants, rs4452212 and rs1599750, not previously associated with ancestry inference in literature, were selected specifically for their capability to differentiate the Middle Eastern cluster. Information on these variants can be found in Figure 2 which also contains plots of their allele frequencies within the cluster used for calculating δ, as well as their frequencies within subpopulations of an independent Middle Eastern dataset [38]. The rs4452212 marker was chosen as a strong candidate to add differentiating power for Middle Eastern versus European clusters. This intergenic variant close to UBBP1 has previously been linked to telomere length variation within Europe, as well as between Europe and Africa [71]. The second marker, rs1599750, supplements and balances the separation for Middle Eastern versus South Asian cluster differentiation. This intergenic variant close to CTDSP2 has previously been reported to show strong geographical genetic variation among the Spanish (CAT), Yoruba (YRI), European (CEU), and Asian (CHB and JPT) populations in a study of microRNA genomic regions in human disease [72].

**Figure 2.**
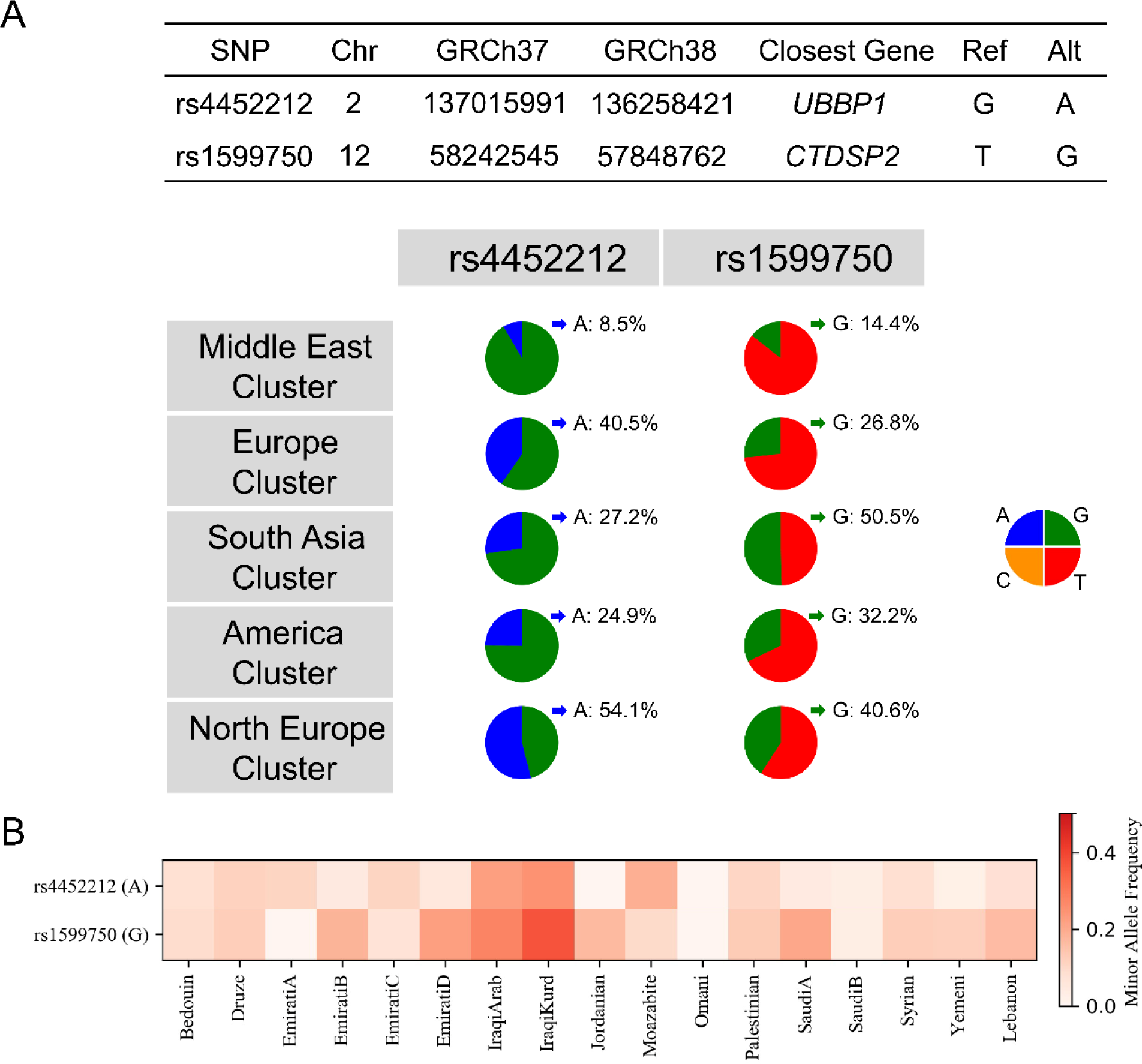
Candidate ME AIM variant information. A) Genomic information and minor allele frequencies of the k means clusters used in marker discovery, and B) Minor allele heat map of an independent dataset of Middle East subpopulations.

An exploration of both variants and their distribution on a geographical scale can be seen in Figure 3, highlighting several proximal subpopulations to the Middle Eastern region. The observed differences between the Middle East and surrounding populations lend support for these markers as candidate ME-AIM variants. However, testing their performance alongside currently known AIM panels is needed to confirm their potential assistance in current BGA inference of individuals from the Middle East.

**Figure 3.**
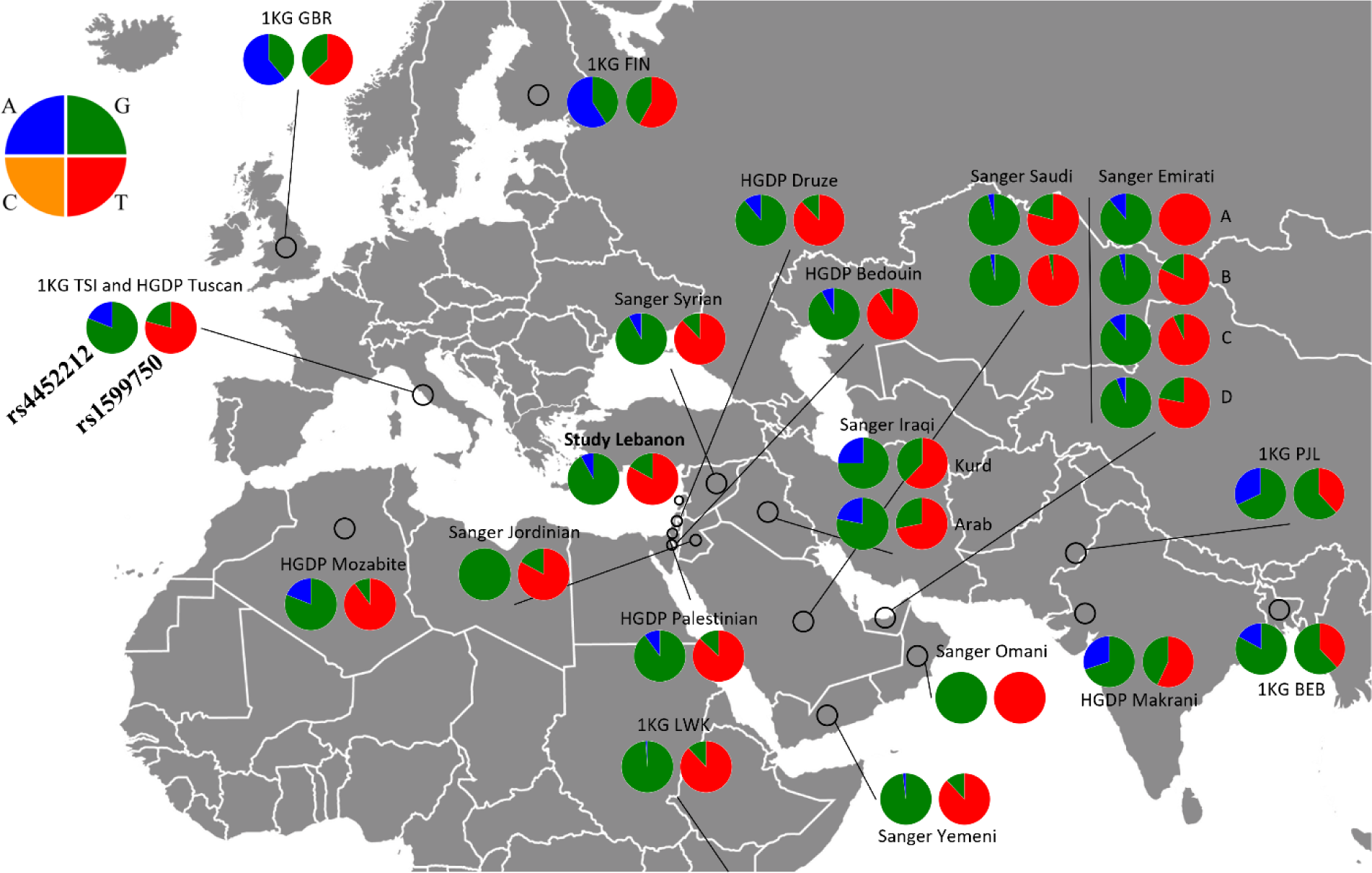
The allele frequencies of rs4452212 and rs1599750 across geographic population data to illustrate its distribution in proximal regions.

### 3.2 Unbiased ancestry assignment label for training and test data using unsupervised clustering

Three whole-genome sequence (WGS) datasets including 1KG, HGDP, and SangerME were combined (n=3,469) and filtered from approximately 180 million to less than 2 million variants in linkage equilibrium (r2 < 0.1) to perform a large ADMIXTURE analysis. This was done in order to ascertain a more complete genetic ancestry classification for both the training and test data used in AIM panel performance assessments. Although K8 provided the lowest cross-validation error (Supp. Fig. 5), we explored the K7 model to observe any key differences (Supp. Fig. 6). The most notable being the split of the East Asian cluster moving from K7 to K8. Notably, this split can be attributed to an East to West axis. This shift in sample assignment can also be seen in a Sankey diagram (Supp Fig. 7). The K8 model was used in further analyses since both models were quite comparable for the majority of the clusters formed. Figure 4A provides an ADMIXTURE plot of K8 for all individuals tested (both training and testing sets), in addition to a filtered training dataset where only samples from 1KG and HGDP that presented proportions at or above the average main ancestry component for the best K were kept as Anchor reference samples (Figure 4B). A breakdown of the distribution of the Anchor reference samples and their assigned clusters can be seen in Figure 4C and 4D, respectively. A list of the samples used for each cluster can be found in Supplemental Table 1.

**Figure 4.**
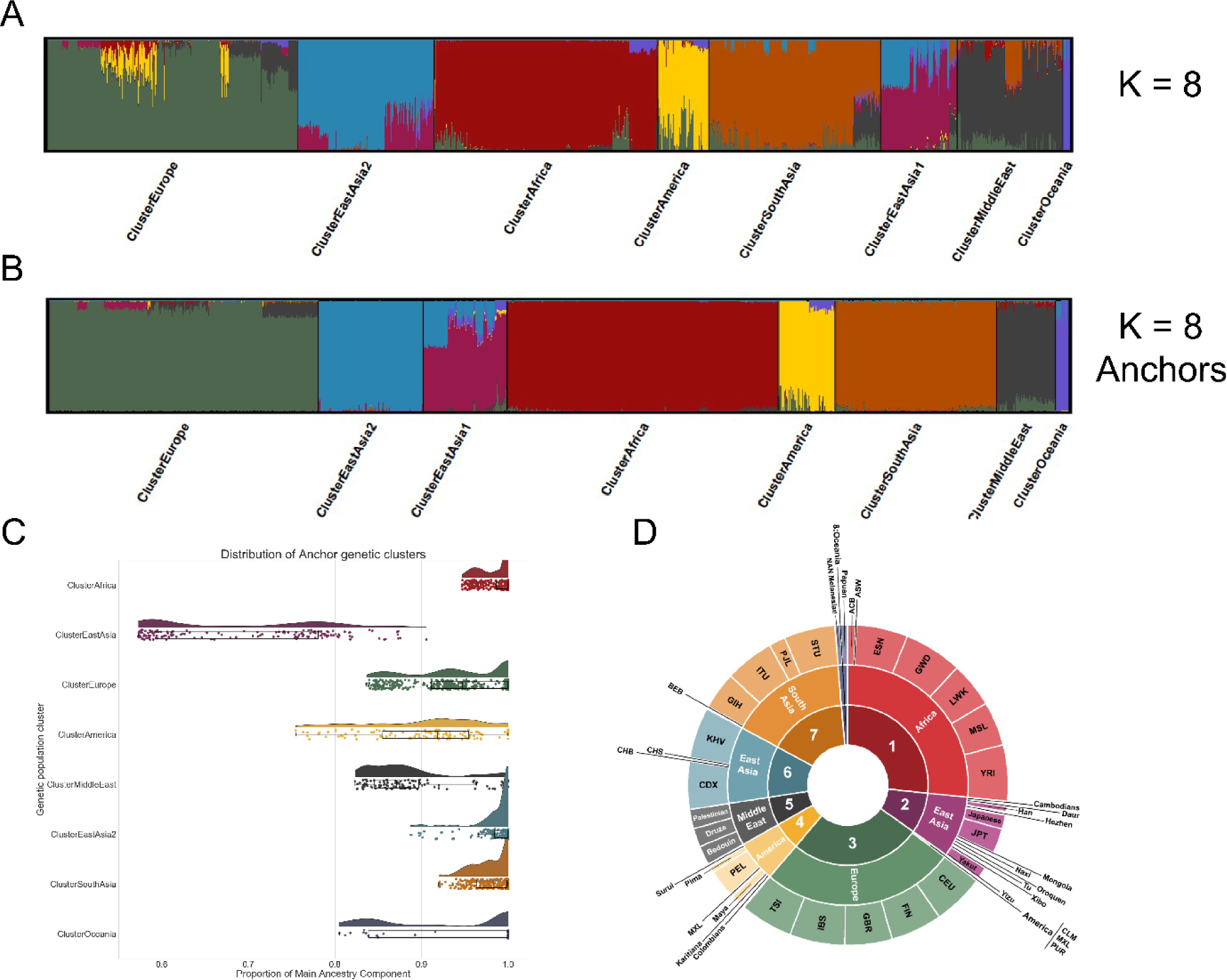
Unsupervised worldwide deep ADMIXTURE of 1KG, HGDP, and SangerME data A) clustered based on maximum K ancestry component per individual and B) SangerME removed and reserved for a “ground truth” validation set and worldwide training data from 1KG and HGDP filtered for samples above the main, mean ancestry component plotted with CLUMPAK. C) Raincloud plot to illustrate the distribution of samples and their main ancestry component within each population cluster with D) membership to each cluster described with pre-defined population and subpopulation information.

The test individuals from the SangerME dataset were assigned to a cluster based on their highest cluster proportion (average ME proportion = 85.5%). All but one (136/137) of the samples clustered with the Middle East cluster at K8. Sample APPG7555904 gave a maximum ancestry component for the Africa cluster followed by the ME cluster (AFR = 0.567; ME = 0.286). Therefore, an AFR assignment was used to simplify and represent the sample’s ancestry designation, or ‘ground truth’, for BGA inference panel testing.

### 3.3. BGA inference panel testing

We assessed the performance of ME BGA inference by compiling the genotypic information from candidate ME-AIMs of this study and from a robust AIM panel described in 3.2. The compiled AIMs were obtained from the Anchor training populations of 1KG and HGDP, in addition to the independent SangerME test dataset. The VISAGE Enhanced Tool (ET) is the latest MPS-based forensic assay for ancestry inference and externally visible trait prediction [18]. Its autosomal AIM panel was compiled for worldwide BGA inference and developed with Middle East differentiation as a primary objective [18]. Although the original panel consisted of 104 variants, two of the autosomal AIMs; rs3857620 and rs17287498, were not included in this study. The former SNP was recently reported to be an uninformative BGA SNP [13, 18] due to inaccurate allele frequency data from the main 1KG Phase III data portal of Ensembl. The latter tri-allelic SNP, was also flagged by Ensembl^79^ for observed ambiguity in alleles. Therefore, the ET panel for this study comparison shall consist of 102 variants. To assess the addition of our two candidate ME markers with the ET panel, it will be referred to as ET+2 for all consecutive analyses. In an effort to include our Middle Eastern individuals from Lebanon in BGA panel assessment, we imputed our Lebanese microarray data using the TOPMED imputation server [73]. However, as 23 of the 102 markers from the Enhanced Tool for comparison were still missing after imputation, we excluded our Lebanese data from the assessments of ET and ET+2.

#### 3.3.1. Snipper and STRUCTURE classification

Snipper [10, 45], the naïve Bayesian classifier, was used to predict the ancestry of the SangerME dataset in relation to the large ADMIXTURE analyses of section 3.2, where 136 out of the 137 were confirmed to have the highest proportion of Middle Eastern ancestry at K8. This allowed us to evaluate the added performance of the two candidate ME-AIMs included from this study (known as ET+2) versus the performance of ET alone for ME-specific BGA inference. Using the Anchor populations as training data which represented eight regions (Supp. Table 1) of AFR, EUR, EAS1, EAS2, ME, SAS, AMR, OCE, both sets of AIMs (ET and ET+2) were able to accurately predict 131 of the 137 SangerME Middle East samples. The sample APPG7555904, which had provided the highest African proportion at K8, was also classified as Middle East for both panels. For most of the misclassified samples (4 out of 6), both ET and ET+2 performed similarly with the same incorrect classifications of three samples with South Asian Ancestry (Table 1) and one sample with a strong likelihood of African ancestry. The two remaining misclassified samples presented contrasting classifications between ET and ET+2, where each panel improved the prediction to Middle East, but overall, a similar accuracy of 95.6% for Middle East inference performed by Snipper was observed using the SangerME test dataset. Although, there was no significant improvement by the inclusion of the two candidate ME-AIMs from this study using the Snipper classifier method, it is important to note that the optimized Anchor populations of K8 (used for training the classifier) may have aided in the accuracy of the model as it provided clearer population boundaries for the snipper likelihood classification.

This was further examined in the STRUCTURE analyses when comparing ET and ET+2 versus the published STRUCTURE proportions from the VISAGE ET data [18], albeit with a K6 model that focused on the use of Eurasian sub-continental reference sets. Evaluations of STRUCTURE and its assignments using ancestry components is an established supplement to any population genetic study and has shown considerable success with relatively small sets of AIMs [11, 19]. When comparing the SangerME dataset using ancestry components derived from the use of the ET and ET+2 AIM panels (Figure 5) to the STRUCTURE proportions of the ET data, there was slight improvement in Middle Eastern ancestry proportions obtained in 7% of individuals (n=10), versus a reduced ME ancestry assignment in 2% of individuals (n=3) (Supp. Table 2). This would also support that improved Anchor reference classification within the training data did yield improved ancestry proportion assignment for the ME region. There were nine incorrect classifications from the SangerME dataset across all comparisons using the highest ancestry proportion as a simplified ancestry assignment. The K6 ET [18] was correct in 117/137 (85%). The K8 ET from this study classified 126/137 (92%) correctly. Lastly, the K8 ET+2 from this study resulted in 125/137 (91%) correctly classified. All of the classification assignments were compared against the ‘ground truth’ highest ancestry proportion assignment. The slight difference between ET versus ET+2 was from the individual previously classified as AFR with highest proportion using the large ADMIXTURE analyses, providing a higher proportion for the ME cluster in ET+2. Another instance of two samples with contrasting correct classifications between ET and ET+2 also occurred, with sample APPG7555894 shifting from SAS to ME and APPG7623338 incorrectly shifting from ME to EUR for ET+2, but overall, there was negligible improvement.

**Figure 5.**
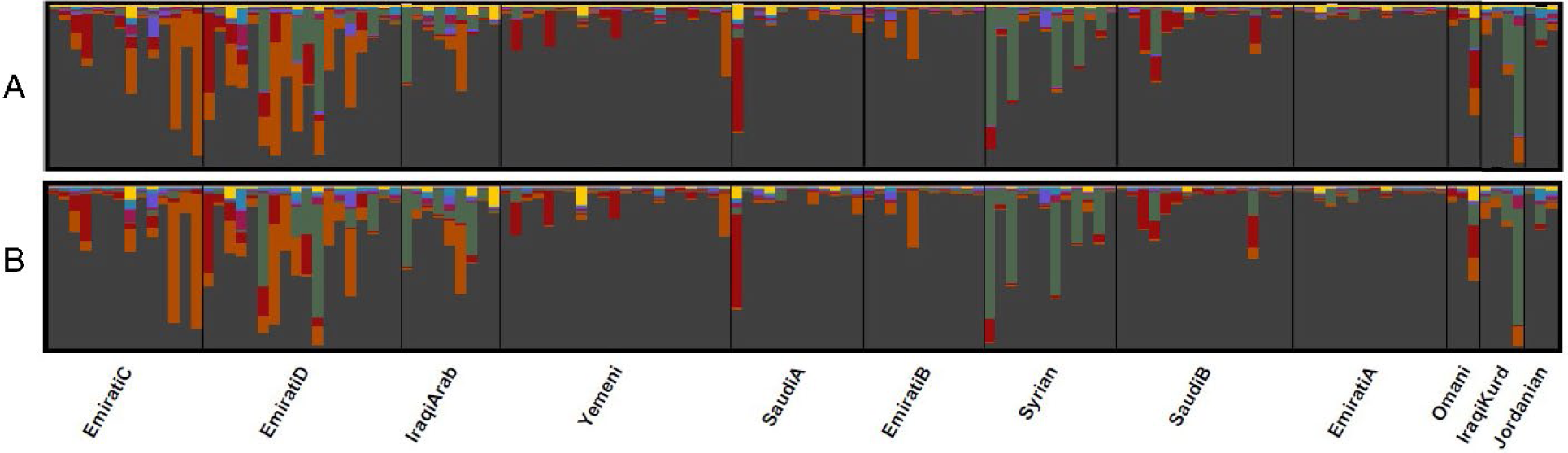
CLUMPAK plots of STRUCTURE analyses for the SangerME samples using the AIMs from A) ET and B) ET+2. See color legend from Figure 4.

#### 3.3.2 Visualizing genetic variance

To explore how best to visualize ancestry and how populations cluster using genetic data as opposed to quantifying or predicting ancestry in the last section, we compared PCA to VAE (popVAE) visualizations using the ET+2 AIM panel (Figure 6). While PCA only captures a combined 35.7% of genetic variance in the first two dimensions, the VAE visualized 100% of the genetic variance in two dimensions alone. This is extremely advantageous for genetic variance interpretability on a 2D scale. Battey et al. [53] suggested that multiple runs with different seeds be conducted for heuristic examination of learning curves to find models with good fit, rather than overfit or underfit [53]. Using popVAE [53] and the Anchor reference data, we found a model for ET+2 that presented a very good fit of validation and training curves (Supp. Fig. 8). In order to optimize the space even further, we ran the model with double the epochs (n=500) and triple the patience (n=150) which provided us with a trained stable reference ancestry space. These tuned hyperparameters simply increased the number of iterations and amount of time allotted for the VAE model to train and validate. For application, the default model architecture of this tool was mostly preserved, but it was necessary to supplement popVAE with in-house scripts to retain the stable and fixed ancestry reference latent space for unknown sample projections. We have developed this model as a web tool, so that other investigators can project individuals of unknown ancestry into the ancestry space of these trained regions/samples. More information can be found on our website at https://walshlab.indianapolis.iu.edu/ under genetic tools. Although PCA also caters for unknown sample projection into a fixed ancestry space, it does not fully reduce the complete genetic variance of the samples into two dimensions and is subject to subtle shifts in coordinates when plotting PCA scatterplots if sample projection into a fixed ancestry space is not implemented.

**Figure 6.**
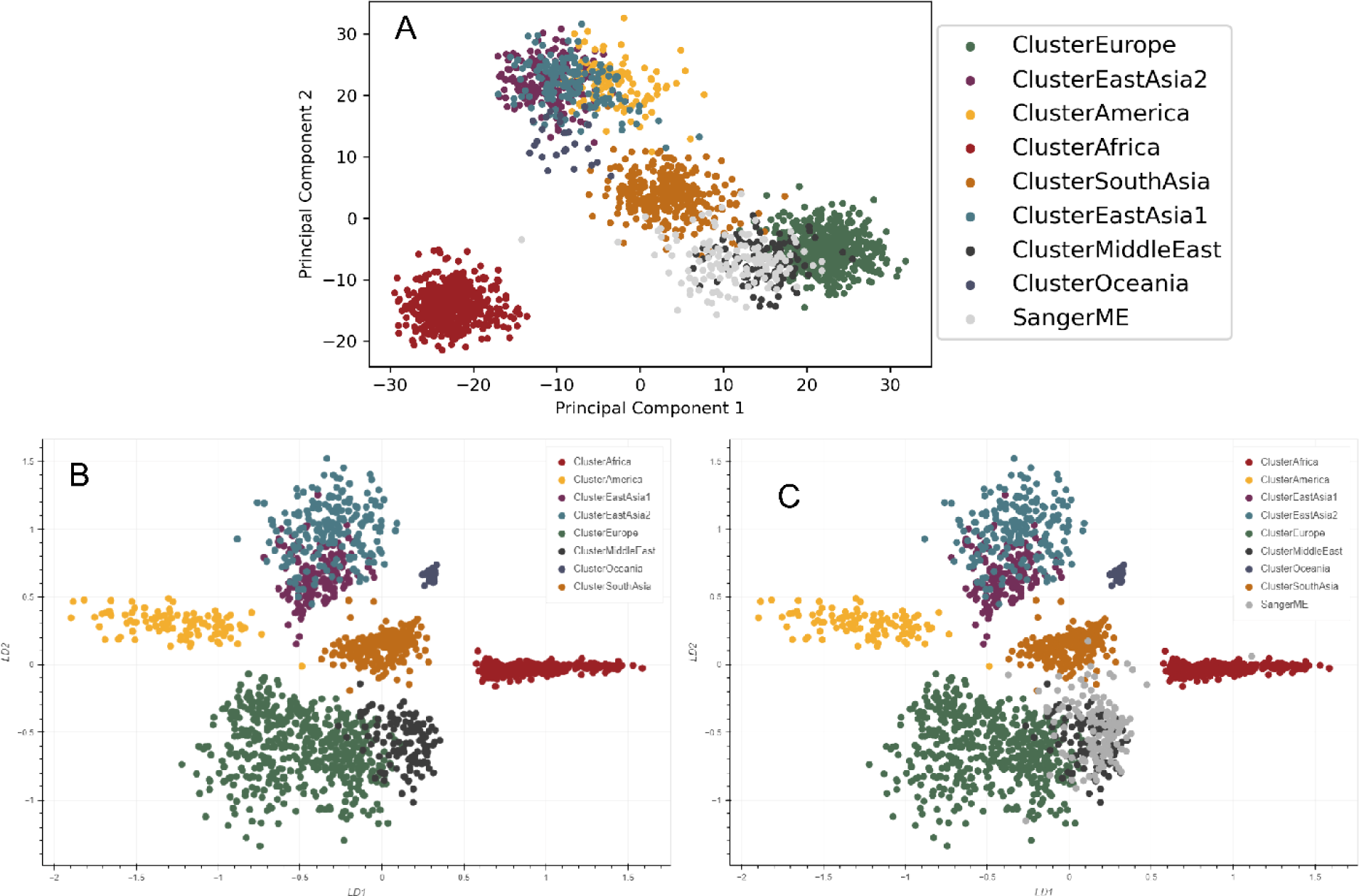
Visualizing genetic variance with worldwide Anchor reference data using A) PCA plot of ET+2 with SangerME test data using the first 2 PC dimensions, B) popVAE plot of latent ancestry reference space, and C) popVAE with projection of SangerME test data samples onto the latent ancestry space.

The test samples from SangerME (n=137), also referred to as our validation samples, were projected onto the two-dimensional ancestry space built using our worldwide Anchor reference population clusters. Out of the 137 individuals, 117 were plotted amongst the Middle Eastern Anchor training data, resulting in 86% accuracy compared to the ground truth admixture results where only one sample was not inferred to represent Middle Eastern ancestry. Population centroids and each test samples’ latent coordinates were used to calculate Euclidean distance and determine which population centroid was closest to each sample (Supp. Table 3). Samples APPG7555904 and APPG7836404, which presented with the highest African proportion at K:8 and were predicted for African ancestry by ET and ET+2, respectively, did plot closer to the African centroid. Sample APPG7555947 plotted closest to the European centroid, also in accordance with its largest ancestry component represented by Europe in both ET and ET+2. However, there were 17 individuals that were closest to the South Asian centroid, with only 4 of those also predicted for South Asian ancestry using ET and ET+2. The remaining 13 individuals closest to the South Asian centroid contrasted with the ET and ET+2 predictions as 11 were predicted Middle Eastern ancestry and 2 were predicted European ancestry.

Additional samples [18] (n=368) with greater than 90% genotype coverage were also selected and uploaded to illustrate the general performance of the web tool in other regions of the world as can be seen in Supplemental Figure 10 and Supplemental Table 4. The provided labels from Ruiz-Ramírez et al. (2023)^18^ have been included in this comparison, which represent several in-house samples as well as individuals from the Simons Genome Diversity Project (SGDP) [74]. With regards the individuals labelled as Middle Eastern, 56 of the 59 (95%) individuals were most proximal to the Middle Eastern centroid. The three misclassified samples fell closer to either European or South Asian centroids. All 29 individuals classified North African fell between the Middle Eastern and African centroids although more proximal to the former. East African individuals shared a similar pattern, however, 21 of the 47 individuals did fall closer to the African Centroid. Only 2 of the 12 individuals with African labels had misclassified shorter distances to the Middle Eastern population centroid however their Population labels indicated a North African region. East Asian individuals presented 28 of 35 matches, 7 of which were found closest to population centroids of South Asia, America, and Europe. Oceanian individuals reported 26 matching, 15 East Asian, and 7 South Asian classifications. There were 16 of 19 correct inferences of South Asian ancestry, with the remaining three falling closer to the East Asian, Oceanian, or Middle Eastern centroids. For the individuals representing Europe, 21 of the 74 were closest to the European centroid. Those mismatched inferences fell closer to population centroids representing the Middle East (49), South Asia (3), and America (1) which align more closely with West Eurasian regions when looking at their Population labels. Lastly, all 10 American individuals were concordant, and the Admixed American individuals had closest centroid distances to Africa (15), Middle East (15), Europe (4), and South Asia (1). These results all indicate there is room for improvement in the development of the ancestry space to allow more levels of BGA inference for individuals, particularly for individuals with higher levels of admixture. It is also worth noting that all of these samples were missing (e.g. NN) the two additional AIM genotypes representing the ET+2 that have been highlighted in this study and may help resolve the difficulty experienced when inferring the European individuals. Overall, this additional assessment was meant to demonstrate the global utility of this ancestry visualization web tool even with small levels (up to 10%) of missing genotypic data.

## 4. Discussion

Recent literature has described the challenges of Middle Eastern population differentiation stemming from its geographical proximity to several continental regions and its observed gene flow throughout history [22, 33, 35]. With these challenges in mind, we opted to compare the most genetically similar clusters with the Middle East (Figure 1C) to find candidate AIMs that may assist in further resolving ancestry inference in the region. The increase in ME sample size and allelic frequency differences (delta; δ) of the Lebanese population assisted in δ score comparisons for the ME region and led us to identify two candidate ME-AIMs. Although we included novel population data from Lebanon in the AIM discovery process for the Middle East, it is worthy to note the limitations of using SNP array data. Even though we were able to improve Middle East sample size from HGDP by over two-fold, our SNP count was restricted to less than one million markers. Nonetheless, the Illumina MEGA microarray has not been used in previous AIMs discovery studies for the Middle East and did provide two candidate AIMs when using overlapping 1KG and HGDP WGS data. Although less cost effective, WGS data is suggested for broader AIMs discovery in the future, whether studies are focused on the ME region or beyond.

With regards the development of Anchor reference datasets for training, many individuals were given memberships in genetic clusters representative of their pre-defined geographical label. However, it is worthy to note that this was not always the case and there were several reference samples grouped into genetic clusters represented by another major population. This occurred much more frequently in samples presenting higher levels of admixture (e.g. individuals with a pre-defined American label grouped with the European Cluster). Therefore, we opted to maximize the cluster separation by utilizing Anchor populations that grouped according to the unsupervised ADMIXTURE clustering method, before pruning according to their mean highest ancestry proportion. This allowed us to still retain core cluster information with real world ancestry proportions. We suggest that ancestry inference of an unknown may be better informed by viewing what reference samples (and their original geographical label according to 1KG and HGDP) were within the population cluster they have been assigned (Figure 4D) to have a better understanding of the labels associated with the individuals with which they cluster.

Incorporating the two candidate ME aims identified in this study, while not resulting in a statistically significant enhancement in ME ancestry classification compared to the utilization of the VISAGE autosomal ET AIMs [18], does contribute substantively to elevated ME ancestry proportions in specific individuals (Supp. Table 2). Consequently, the integration of these markers in future designs is potentially advantageous, especially considering their presence in various SNP array designs, facilitating seamless implementation. Furthermore, these markers may serve as viable substitutes for variants within the ET panel that do not amplify adequately or experience dropout.

PCA has been a standard in the field to illustrate population clustering for the last decade. However, recent literature has shed light on its biased nature and inability to be reliable, robust, or repeatable [52]. Additionally, due to the nature of PCA, it is necessary to visualize multiple dimensions at once to observe adequate cluster separation (i.e., first, second, third, and fourth dimensions at minimum). In an effort to simplify the visualization space and maximize the observable reduced genetic variance for population clusters into less dimensions, we explored the popVAE variational autoencoder with three proponents in mind: First, it can incorporate non-linear relationships representative of population genetic data, second, it allows users to define the dimensionality of the latent space, which for the sake of easy visualization, this is best represented in a 2D (two latent dimensions) space, and third, it has the ability to preserve global geometry which is defined as the distance among clusters equating to geography or genetic differentiation [53].

The application of a VAE as an alternative approach to best capture genetic variance could provide the community with better regional differentiation capabilities. Here, we first tested this deep learning approach on AIMs which resulted in highly distinct population clustering amongst 100% of the genetic variance in two dimensions. The majority of the test samples from the SangerME dataset clustered well with trained ME clusters providing strong support for the use of Autoencoders to correctly distinguish clusters using genetic data with adequate training epochs and offering a standardized and stable ancestry space. In comparison to the PCA on the first two dimensions, where there is significant overlap in the East Asian and American clusters, there is clear separation for all clusters using the AE-built reference space which provides greater resolution for unknown samples. With regards to admixture, further testing of this ancestry space will shed light on how individuals cluster, however, it is expected that they would fall within or close to the cluster representing their highest ancestry proportion. In practice, we observed that over 90% of the Middle Eastern test samples were proximal to their population centroid in reference to the ET and ET+2 STRUCTURE outcomes, and misclassifications represented admixed individuals plotting in between two population centroids, namely Middle Eastern and South Asian centroids, though slightly closer to the latter.

We encourage the further use and testing of this approach using our freely available web tool for the projection of samples with AIM genotypes onto our trained ancestry reference space described here. While the use of variational autoencoders for ancestry visualization certainly holds promise, we also recommend exploring alternative autoencoder models in population genetics [75], with a specific focus on their effectiveness in ancestry inference utilizing AIMs. We also recognize and encourage existing alternative approaches for marker discovery and inference evaluation that include unsupervised methods [44], supervised methods [76], machine learning models [77], deep learning models [78], and multivariate statistics [36]. These techniques may greatly benefit the community, particularly as we curate the best combination of ancestry informative markers and integrate multi-tier stepwise approaches with deep-learning applications to best visualize and ideally predict unknown individuals from trained and fixed ancestry spaces for standardized and replicable ancestry inference.

## 5. Conclusion

The unique geographical position and cultural history of the Middle Eastern region pose persistent challenges in BGA inference, especially in the context of using AIMs panels. We explored Middle Eastern ancestry inference with the addition of Middle Eastern population data from Lebanon and in the context of genetic clustering in lieu of geographical labels. Two candidate AIMs were reported and evaluated alongside the VISAGE Enhanced Tool using Snipper, STRUCTURE, PCA, and popVAE. We report a worldwide deep ADMIXTURE analysis on WGS data and provide stable ancestry components for samples within 1KG, HGDP, and SangerME for reference of their genetic ‘ground truth’. Lastly, we propose the utilization of deep learning approaches such as autoencoders for building two-dimensional reference ancestry latent spaces to be used in future visualizations of population variance using AIMs for forensic ancestry inference. We also provide an open-source web tool for investigators to plot unknown (or known) samples into a stable latent ancestry space. Our findings not only contribute to overcoming challenges in Middle Eastern BGA inference with a forensic application but also pave the way for more robust and nuanced methodologies in ancestry inference as a whole.

## Acknowledgements

We acknowledge and thank all the individuals that participated in these studies, without them this research would not be possible. For the usage of Indiana University servers to perform computational work, this research was supported in part by Lilly Endowment, Inc., through its support for the Indiana University Pervasive Technology Institute and by the Indiana Genomics Initiative. IUI personnel, data collection, and analyses were supported in part by the US National Institute of Justice (2014-DN-BX-K031).

## CRediT authorship contribution statement

**Noah Herrick**: Writing – original draft, Writing – review & editing, Investigation, Methodology, Software, Visualization, Conceptualization. **Mirna Ghemrawi**: Resources, Writing – review & editing. **Sylvia Singh**: Visualization, Writing – review & editing. **Rami Mahfouz**: Resources, Writing – review & editing. **Susan Walsh**: Writing – original draft, Writing – review & editing, Investigation, Resources, Conceptualization.

## References

1. Walsh, S., et al., IrisPlex: A sensitive DNA tool for accurate prediction of blue and brown eye colour in the absence of ancestry information. Forensic Science International: Genetics, 2011. 5(3): p. 170–180.

2. Walsh, S., et al., Developmental validation of the HIrisPlex system: DNA-based eye and hair colour prediction for forensic and anthropological usage. Forensic Science International: Genetics, 2014. 9: p. 150–161.

3. Chaitanya, L., et al., Bringing colour back after 70 years: Predicting eye and hair colour from skeletal remains of World War II victims using the HIrisPlex system. Forensic Science International: Genetics, 2017. 26: p. 48–57.

4. Walsh, S., et al., Global skin colour prediction from DNA. Human Genetics, 2017. 136(7): p. 847–863.

5. Pośpiech, E., et al., Towards broadening Forensic DNA Phenotyping beyond pigmentation: Improving the prediction of head hair shape from DNA. Forensic Science International: Genetics, 2018. 37: p. 241–251.

6. Chaitanya, L., et al., The HIrisPlex-S system for eye, hair and skin colour prediction from DNA: Introduction and forensic developmental validation. Forensic Science International: Genetics, 2018. 35: p. 123–135.

7. Shriver, M.D., et al., Ethnic-affiliation estimation by use of population-specific DNA markers. American journal of human genetics, 1997. 60(4): p. 957.

8. Rosenberg, N.A., et al., Informativeness of genetic markers for inference of ancestry. The American Journal of Human Genetics, 2003. 73(6): p. 1402–1422.

9. Frudakis, T., et al., A classifier for the SNP-based inference of ancestry. Journal of forensic sciences, 2003. 48(4): p. 771–782.

10. Phillips, C., et al., Inferring ancestral origin using a single multiplex assay of ancestry-informative marker SNPs. Forensic Science International: Genetics, 2007. 1(3-4): p. 273–280.

11. Phillips, C., Forensic genetic analysis of bio-geographical ancestry. Forensic Science International: Genetics, 2015. 18: p. 49–65.

12. Schneider, P.M., B. Prainsack, and M. Kayser, The use of forensic DNA phenotyping in predicting appearance and biogeographic ancestry. Deutsches Ärzteblatt International, 2019. 116(51-52): p. 873.

13. Kayser, M., et al., Recent advances in Forensic DNA Phenotyping of appearance, ancestry and age. Forensic Science International: Genetics, 2023: p. 102870.

14. Kling, D., et al., Investigative genetic genealogy: Current methods, knowledge and practice. Forensic Science International: Genetics, 2021. 52: p. 102474.

15. de Vries, J.H., et al., Impact of SNP microarray analysis of compromised DNA on kinship classification success in the context of investigative genetic genealogy. Forensic Science International: Genetics, 2022. 56: p. 102625.

16. Glynn, C.L., Bridging disciplines to form a new one: the emergence of forensic genetic genealogy. Genes, 2022. 13(8): p. 1381.

17. Daeid, N.N., et al., The analysis of ancestry with small-scale forensic panels of genetic markers. Emerging Topics in Life Sciences, 2021. 5(3): p. 443–453.

18. Ruiz-Ramírez, J., et al., Development and evaluations of the ancestry informative markers of the VISAGE Enhanced Tool for Appearance and Ancestry. Forensic Science International: Genetics, 2023. 64: p. 102853.

19. Phillips, C., et al., Building a forensic ancestry panel from the ground up: The EUROFORGEN Global AIM-SNP set. Forensic Science International: Genetics, 2014. 11: p. 13–25.

20. Jäger, A.C., et al., Developmental validation of the MiSeq FGx forensic genomics system for targeted next generation sequencing in forensic DNA casework and database laboratories. Forensic Science International: Genetics, 2017. 28: p. 52–70.

21. Al-Asfi, M., et al., Assessment of the precision ID ancestry panel. International journal of legal medicine, 2018. 132: p. 1581–1594.

22. Pereira, V., et al., Development and validation of the EUROFORGEN NAME (North African and Middle Eastern) ancestry panel. Forensic Science International: Genetics, 2019. 42: p. 260–267.

23. Phillips, C., et al., MAPlex-A massively parallel sequencing ancestry analysis multiplex for Asia-Pacific populations. Forensic Science International: Genetics, 2019. 42: p. 213–226.

24. Phillips, C., et al., A compilation of tri-allelic SNPs from 1000 Genomes and use of the most polymorphic loci for a large-scale human identification panel. Forensic Science International: Genetics, 2020. 46: p. 102232.

25. Diepenbroek, M., et al., Evaluation of the ion Ampliseq™ phenotrivium panel: MPS-based assay for ancestry and phenotype predictions challenged by casework samples. Genes, 2020. 11(12): p. 1398.

26. De la Puente, M., et al., Development and evaluation of the ancestry informative marker panel of the VISAGE basic tool. Genes, 2021. 12(8): p. 1284.

27. Tillmar, A., et al., The FORCE Panel: An all-in-one SNP marker set for confirming investigative genetic genealogy leads and for general forensic applications. Genes, 2021. 12(12): p. 1968.

28. Rauf, S., et al., Unveiling forensically relevant biogeographic, phenotype and Y-chromosome SNP variation in Pakistani ethnic groups using a customized hybridisation enrichment forensic intelligence panel. Plos one, 2022. 17(2): p. e0264125.

29. Cavalli-Sforza, L.L., et al., The history and geography of human genes. 1994: Princeton university press.

30. Grugni, V., et al., Ancient migratory events in the Middle East: new clues from the Y-chromosome variation of modern Iranians. PloS one, 2012. 7(7): p. e41252.

31. Haber, M., et al., Influences of history, geography, and religion on genetic structure: the Maronites in Lebanon. European Journal of Human Genetics, 2011. 19(3): p. 334–340.

32. Haber, M., et al., Genome-wide diversity in the levant reveals recent structuring by culture. PLoS genetics, 2013. 9(2): p. e1003316.

33. Phillips, C., et al., Eurasiaplex: A forensic SNP assay for differentiating European and South Asian ancestries. Forensic Science International: Genetics, 2013. 7(3): p. 359–366.

34. Truelsen, D., et al., Typing of two middle eastern populations with the precision ID ancestry panel. Forensic Science International: Genetics Supplement Series, 2017. 6: p. e301–e302.

35. Truelsen, D., et al., Assessment of the effectiveness of the EUROFORGEN NAME and Precision ID Ancestry panel markers for ancestry investigations. Scientific Reports, 2021. 11(1): p. 18595.

36. Pilli, E., et al., Biogeographical ancestry, variable selection, and PLS-DA method: a new panel to assess ancestry in forensic samples via MPS technology. Forensic Science International: Genetics, 2023. 62: p. 102806.

37. Bergström, A., et al., Insights into human genetic variation and population history from 929 diverse genomes. Science, 2020. 367(6484): p. eaay5012.

38. Almarri, M.A., et al., The genomic history of the Middle East. Cell, 2021. 184(18): p. 4612–4625. e14.

39. Haber, M., et al., Continuity and admixture in the last five millennia of Levantine history from ancient Canaanite and present-day Lebanese genome sequences. The American Journal of Human Genetics, 2017. 101(2): p. 274–282.

40. Wright, S., The genetical structure of populations. Annals of eugenics, 1949. 15(1): p. 323–354.

41. Weir, B.S. and C.C. Cockerham, Estimating F-statistics for the analysis of population structure. evolution, 1984: p. 1358–1370.

42. Alexander, D.H., J. Novembre, and K. Lange, Fast model-based estimation of ancestry in unrelated individuals. Genome Research, 2009. 19(9): p. 1655–1664.

43. Pritchard, J.K., M. Stephens, and P. Donnelly, Inference of Population Structure Using Multilocus Genotype Data. Genetics, 2000. 155(2): p. 945–959.

44. Ko, S., et al., Unsupervised discovery of ancestry-informative markers and genetic admixture proportions in biobank-scale datasets. The American Journal of Human Genetics, 2023.

45. Fondevila, M., et al., Revision of the SNPforID 34-plex forensic ancestry test: assay enhancements, standard reference sample genotypes and extended population studies. Forensic Science International: Genetics, 2013. 7(1): p. 63–74.

46. Menozzi, P., A. Piazza, and L. Cavalli-Sforza, Synthetic Maps of Human Gene Frequencies in Europeans: These maps indicate that early farmers of the Near East spread to all of Europe in the Neolithic. Science, 1978. 201(4358): p. 786-792.

47. Patterson, N., A.L. Price, and D. Reich, Population structure and eigenanalysis. PLoS genetics, 2006. 2(12): p. e190.

48. Lao, O., et al., Correlation between genetic and geographic structure in Europe. Current Biology, 2008. 18(16): p. 1241–1248.

49. Novembre, J., et al., Genes mirror geography within Europe. Nature, 2008. 456(7218): p. 98-101.

50. Novembre, J. and M. Stephens, Interpreting principal component analyses of spatial population genetic variation. Nature genetics, 2008. 40(5): p. 646–649.

51. Tian, C., et al., European population genetic substructure: further definition of ancestry informative markers for distinguishing among diverse European ethnic groups. Molecular Medicine, 2009. 15: p. 371–383.

52. Elhaik, E., Principal component analyses (PCA)-based findings in population genetic studies are highly biased and must be reevaluated. Scientific Reports, 2022. 12(1): p. 14683.

53. Battey, C.J., G.C. Coffing, and A.D. Kern, Visualizing population structure with variational autoencoders. G3 Genes|Genomes|Genetics, 2021. 11(1): p. 1-11.

54. Herrick, N. and S. Walsh, ILIAD: a suite of automated Snakemake workflows for processing genomic data for downstream applications. BMC bioinformatics, 2023. 24(1): p. 424.

55. The 1000 Genomes Project Consortium. A global reference for human genetic variation. Nature, 2015. 526(7571): p. 68-74.

56. 2023., S.T.D.T.h.t.n.n.n.g.T.s.s.c.v.s.A.M., SRA Toolkit Development Team. https://trace.ncbi.nlm.nih.gov/Traces/sra/sra.cgi?view=software. 2022. Accessed 1 March 2023.

57. Li, H., A statistical framework for SNP calling, mutation discovery, association mapping and population genetical parameter estimation from sequencing data. Bioinformatics, 2011. 27(21): p. 2987–2993.

58. Chang, C.C., et al., Second-generation PLINK: rising to the challenge of larger and richer datasets. GigaScience, 2015. 4(1).

59. Purcell, S., et al., PLINK: a tool set for whole-genome association and population-based linkage analyses. The American journal of human genetics, 2007. 81(3): p. 559–575.

60. Pedersen, T.L., Pedersen, T. L. ggforce: Accelerating ‘ggplot2’. R package version 0.4.1. 2022.

61. Kopelman, N.M., et al., Clumpak: a program for identifying clustering modes and packaging population structure inferences across K. Molecular Ecology Resources, 2015. 15(5): p. 1179–1191.

62. Mölder, F., et al., Sustainable data analysis with Snakemake. F1000Research, 2021. 10: p. 33.

63. Byrska-Bishop, M., et al., High coverage whole genome sequencing of the expanded 1000 Genomes Project cohort including 602 trios. 2021, Cold Spring Harbor Laboratory.

64. Brunson JC, R.Q., Brunson JC, Read QD (2023). “ggalluvial: Alluvial Plots in ‘ggplot2’.” R package version 0.12.5, http://corybrunson.github.io/ggalluvial/.

65. Team, R.C., R Core Team (2021). R: A language and environment for statistical computing. R Foundation for Statistical Computing, Vienna, Austria. URL https://www.R-project.org/.

66. H, W., Wickham H (2016). ggplot2: Elegant Graphics for Data Analysis. Springer-Verlag New York. ISBN 978–3-319-24277-4, https://ggplot2.tidyverse.org.

67. Weihs C, L.U., Luebke K, Raabe N (2005). “klaR Analyzing German Business Cycles.” In Baier D, Decker R, Schmidt-Thieme L (eds.), Data Analysis and Decision Support, 335-343., Weihs C, Ligges U, Luebke K, Raabe N (2005). “klaR Analyzing German Business Cycles.” In Baier D, Decker R, Schmidt-Thieme L (eds.), Data Analysis and Decision Support, 335-343.

68. Schauberger P, W.A., Schauberger P, Walker A (2022). openxlsx: Read, Write and Edit xlsx Files. https://ycphs.github.io/openxlsx/index.html, https://github.com/ycphs/openxlsx.

69. Kuhn, M., Kuhn, Max (2008). “Building Predictive Models in R Using the caret Package.” Journal of Statistical Software, 28(5), 1–26. doi:10.18637/jss.

70. Rosenberg, N.A., Genetic Structure of Human Populations. Science, 2002. 298(5602): p. 2381-2385.

71. Hansen, M.E., et al., Shorter telomere length in Europeans than in Africans due to polygenetic adaptation. Human molecular genetics, 2016. 25(11): p. 2324–2330.

72. Muiños-Gimeno, M., et al., Design and evaluation of a panel of single-nucleotide polymorphisms in microRNA genomic regions for association studies in human disease. European Journal of Human Genetics, 2010. 18(2): p. 218–226.

73. Taliun, D., et al., Sequencing of 53,831 diverse genomes from the NHLBI TOPMed Program. Nature, 2021. 590(7845): p. 290-299.

74. Mallick, S., et al., The Simons genome diversity project: 300 genomes from 142 diverse populations. Nature, 2016. 538(7624): p. 201-206.

75. Yuan, M., et al., Hybrid autoencoder with orthogonal latent space for robust population structure inference. Scientific reports, 2023. 13(1): p. 2612.

76. Pfaffelhuber, P., et al., How to choose sets of ancestry informative markers: A supervised feature selection approach. Forensic Science International: Genetics, 2020. 46: p. 102259.

77. van der Gaag, K., et al., The ForAPP: Forensic Ancestry Prediction Pipeline for the interpretation of ancestry informative markers. Forensic Science International: Genetics Supplement Series, 2022. 8: p. 12–14.

78. Qu, Y., D. Tran, and W. Ma, Deep learning approach to biogeographical ancestry inference. Procedia Computer Science, 2019. 159: p. 552–561.

