## Supplemental Figures for "Exploring ancestry inference of the Middle East"

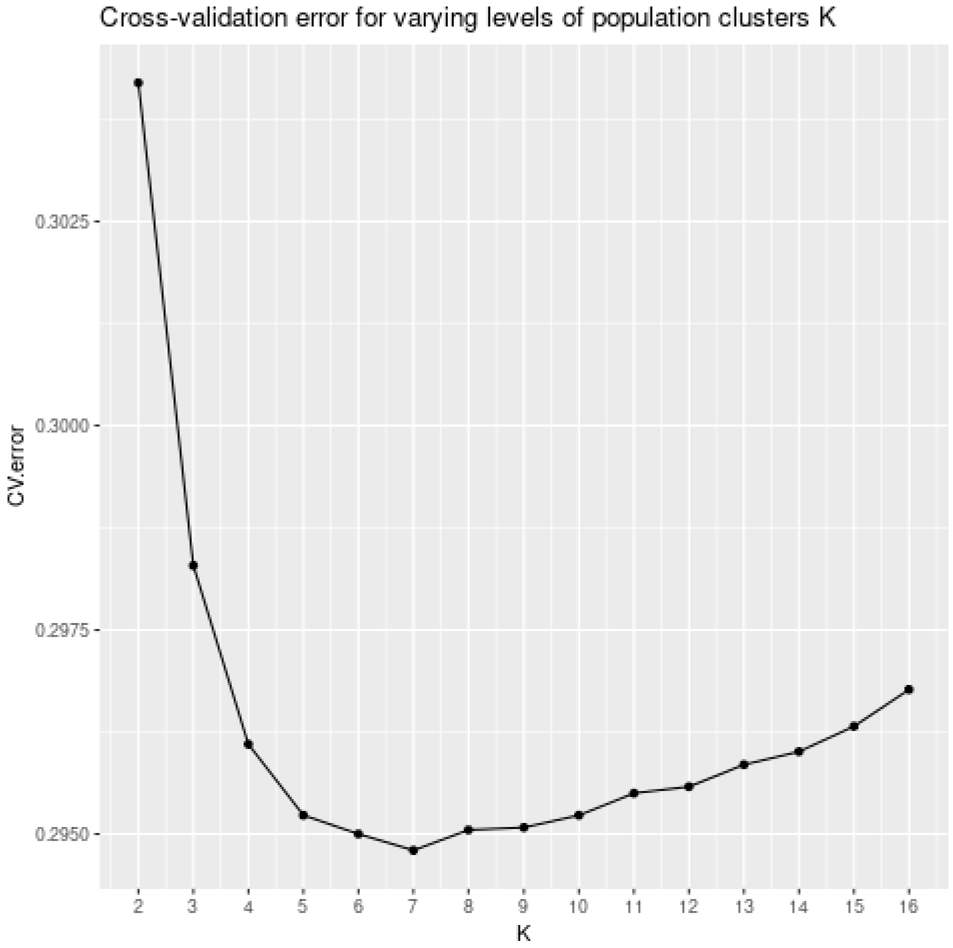


**Supplemental Figure 1** Ten-fold cross-validation errors from unsupervised ADMIXTURE models of K:2-16 for Lebanese, HGDP, and 1KG combined data (samples=2,051; snps=194,321).

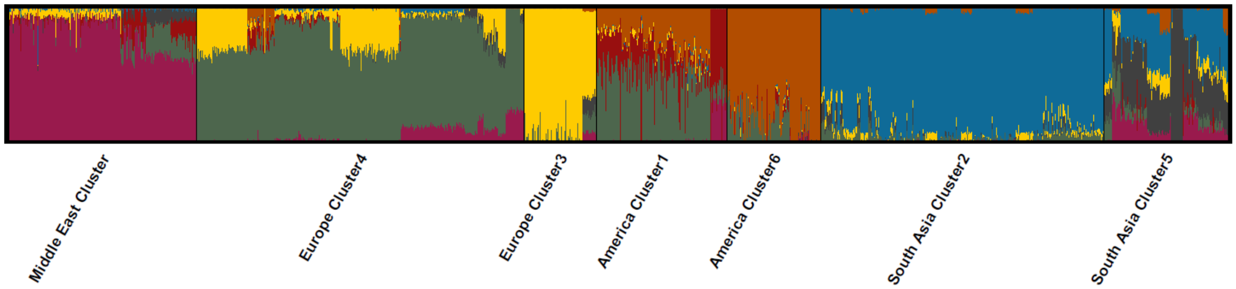


**Supplemental Figure 2** Unsupervised ADMIXTURE model K7 for Lebanese, HGDP, and 1KG combined data. The applied labels coincide with K-means designation of cluster labels based on ancestry components for each sample and represent the unbiased population clusters used in downstream analyses. (SNPs = 194,321; n = 2,051).

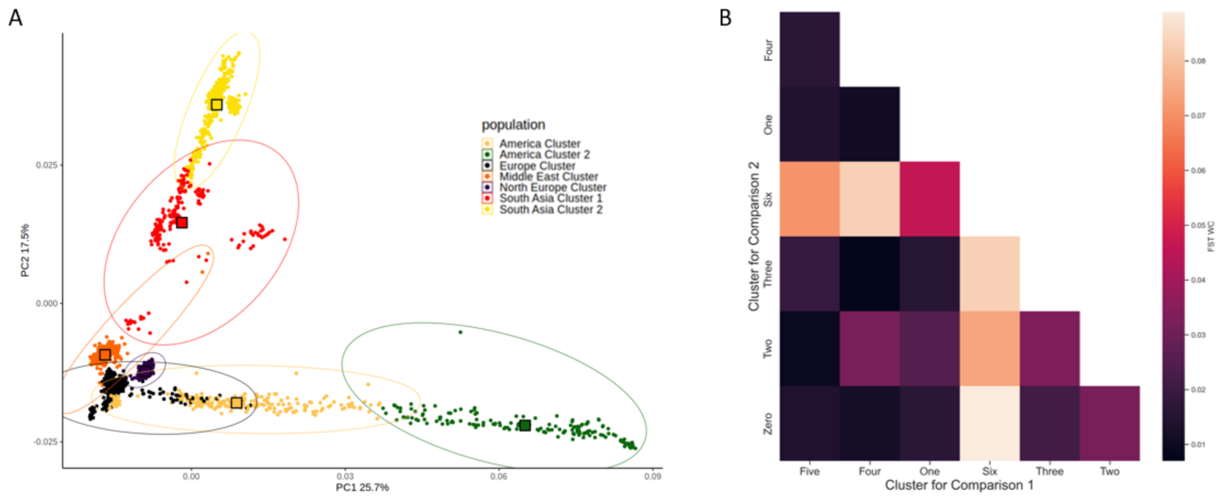


**Supplemental Figure 3** Genetic variance tests to filter for population clusters that were most biogeographically relevant for pairwise cluster comparisons with the Middle East Cluster for AIMs discovery. **A)** PCA of the K7 clusters consisting of Lebanon, HGDP, and 1KG samples. Individuals with East Asian and African ancestries were excluded at this stage for greatest initial observed variance (SNPs = 194,321; n = 2,051). **B)** The same data underwent mean FST pairwise comparisons. This heatmap of the FST results illustrates that Clusters One (America), Three (North Europe), Four (Europe), and Five (South Asia) display less genetic differentiation with the Middle East than do Clusters Two (More distant South Asia Cluster 2 samples) and Clusters Six (More distant America Cluster 2 samples).

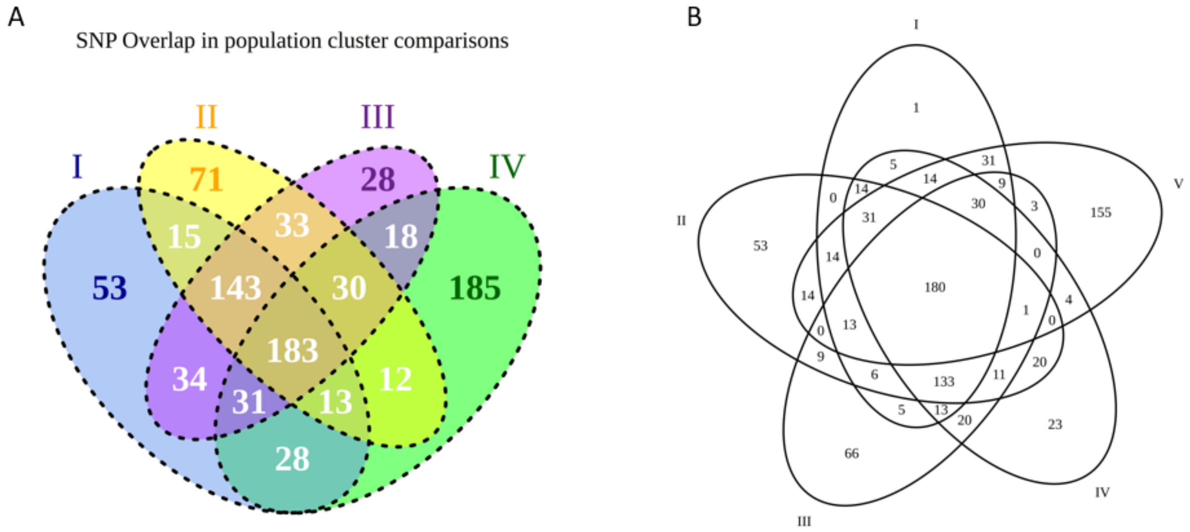


**Supplemental Figure 4** Venn Diagrams representing the selection of markers in the Top 500 **A)** from each of the four comparisons: I) Middle East Cluster vs. America Cluster 1, II) Middle East Cluster vs. North Europe Cluster 3, III) Middle East Cluster vs. Europe Cluster 4, and IV) Middle East Cluster vs. South Asia Cluster 5. The top 500 cumulative markers were not included in this venn diagram for ease of visualization. **B)** Here we include the top 500 cumulative markers. There was only one marker that did not overlap across one of the four other comparisons: I) Cumulative Delta top 500, II) Middle East Cluster vs. America Cluster 1, III) Middle East Cluster vs. North Europe Cluster 3, IV) Middle East Cluster vs. Europe Cluster 4, and V) Middle East Cluster vs. South Asia Cluster 5.

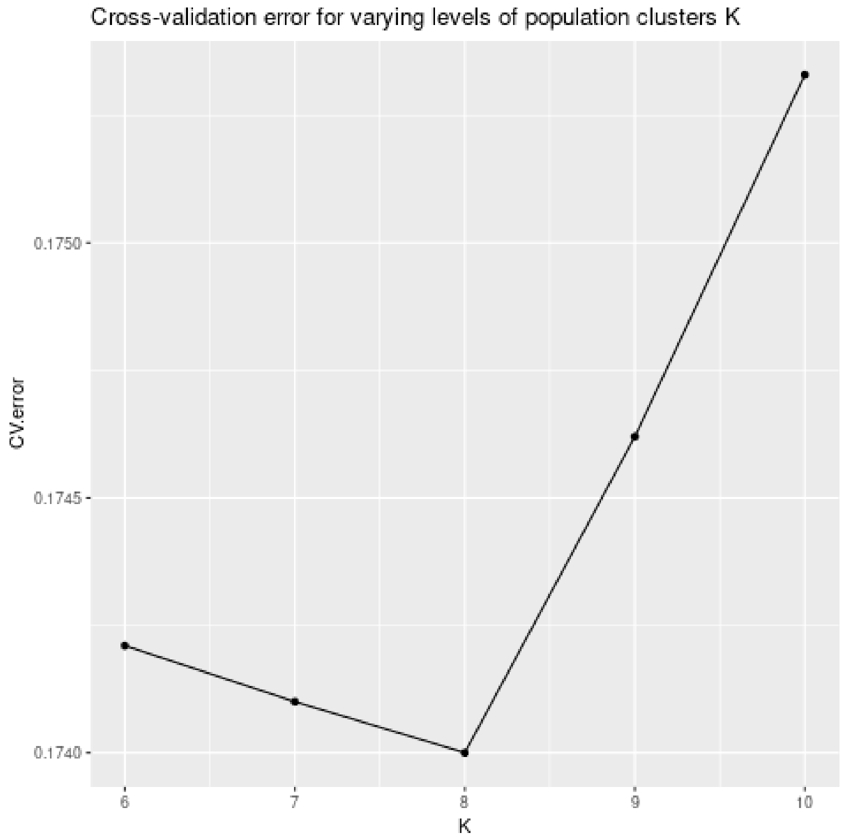


**Supplemental Figure 5** Ten-fold cross-validation errors from unsupervised ADMIXTURE models of K:6-10 for SangerME, HGDP, and 1KG combined data (samples=3,469; snps=1,712,516).

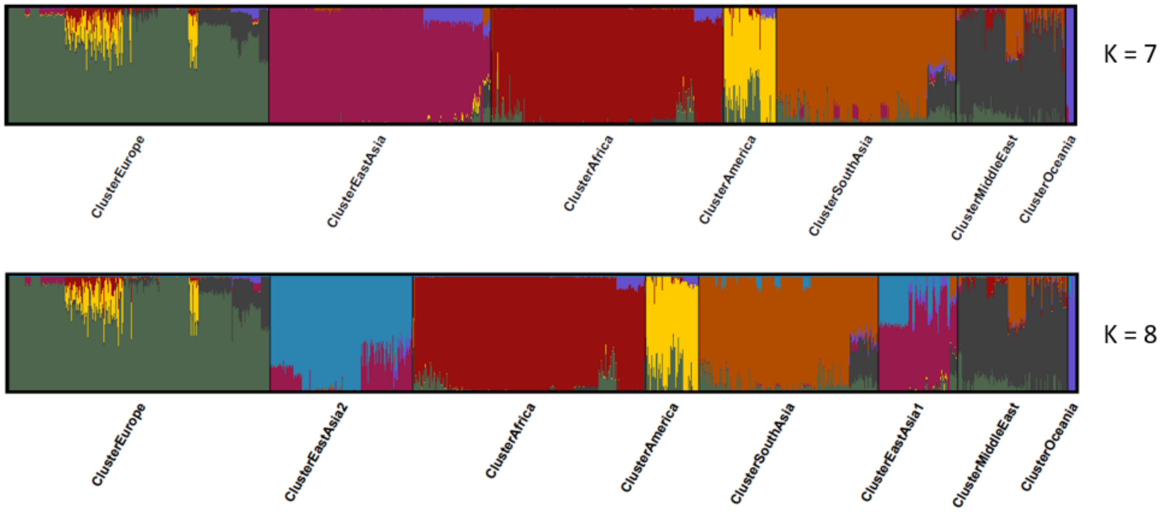


**Supplemental Figure 6** Unsupervised ADMIXTURE model **A)** K7 and **B)** K8 for SangerME, HGDP, and 1KG combined data (snps=1,712,516). The applied labels coincide with maximum K designation of cluster labels based on ancestry components for each sample and represent the unbiased population clusters used in downstream analyses.

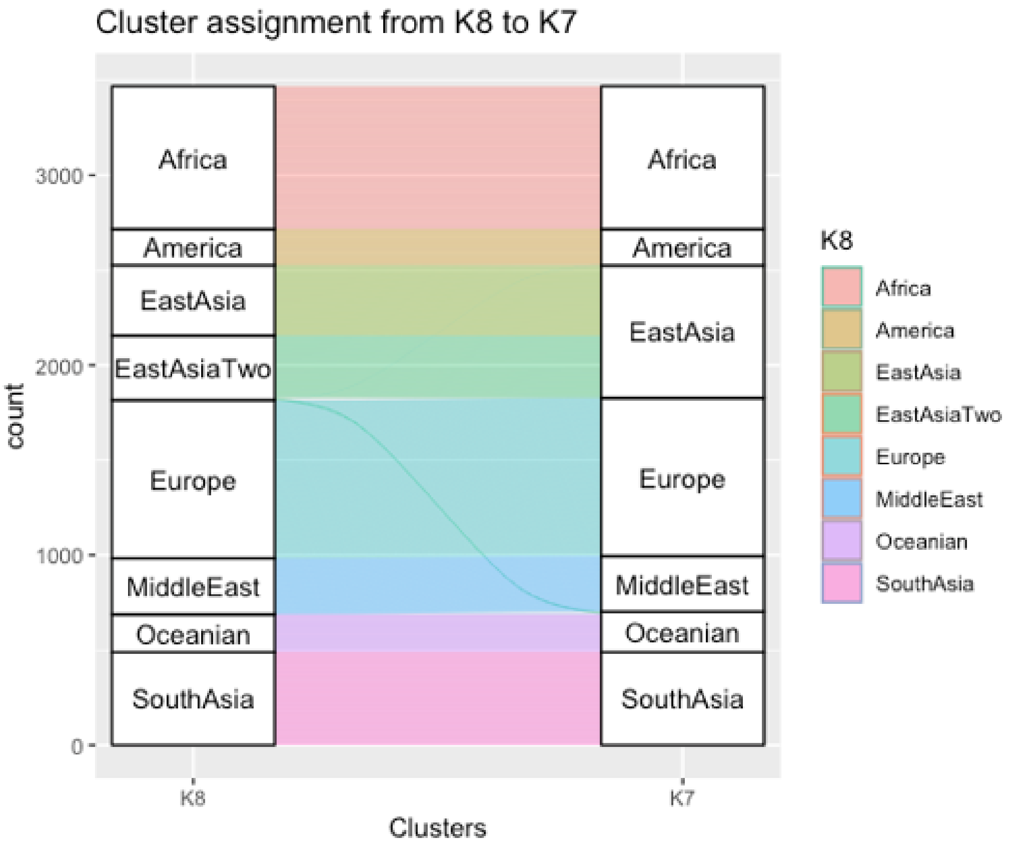


**Supplemental Figure 7** A Sankey plot to visualize the flow of samples observed in our exploration of K:7 and K:8. The East Asia Cluster divides into two distinct population clusters in the flow from K:7 to K:8 (right to left). A number of European Cluster samples that were perhaps more ambiguous at K:7 were assigned to the Europe Cluster at K:8.

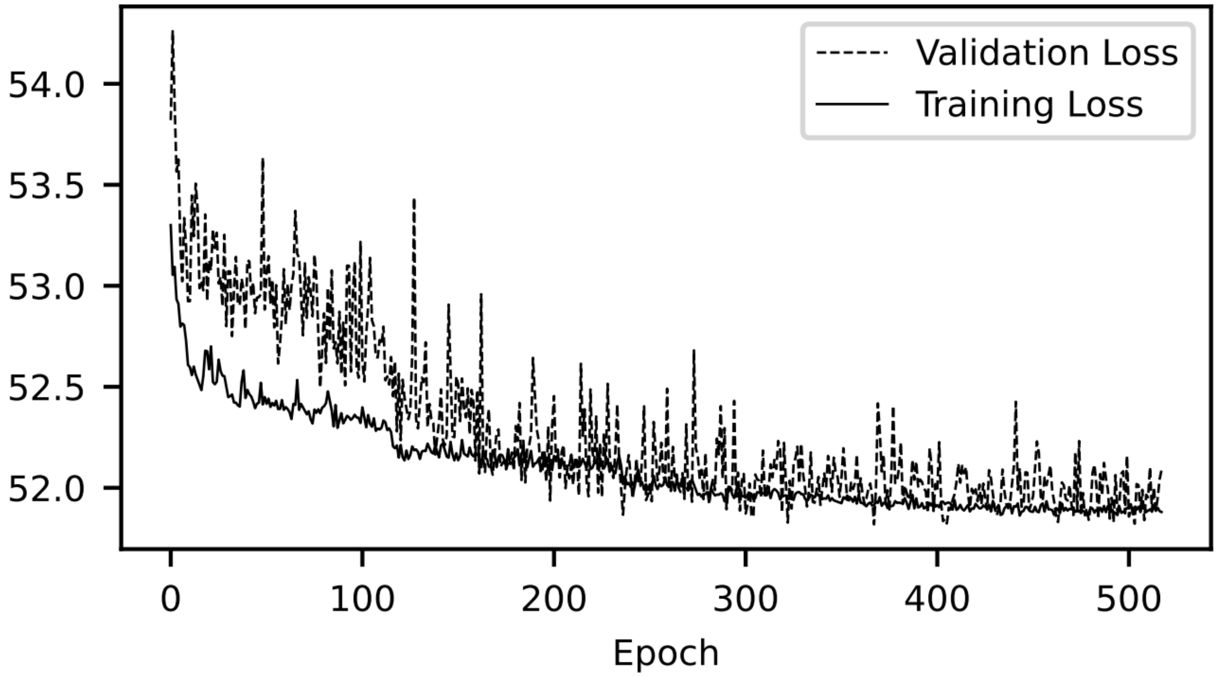


**Supplemental Figure 8** The learning curve associated with the adapted popvae model used for projection of SangerME samples into the Anchor reference ancestry space.

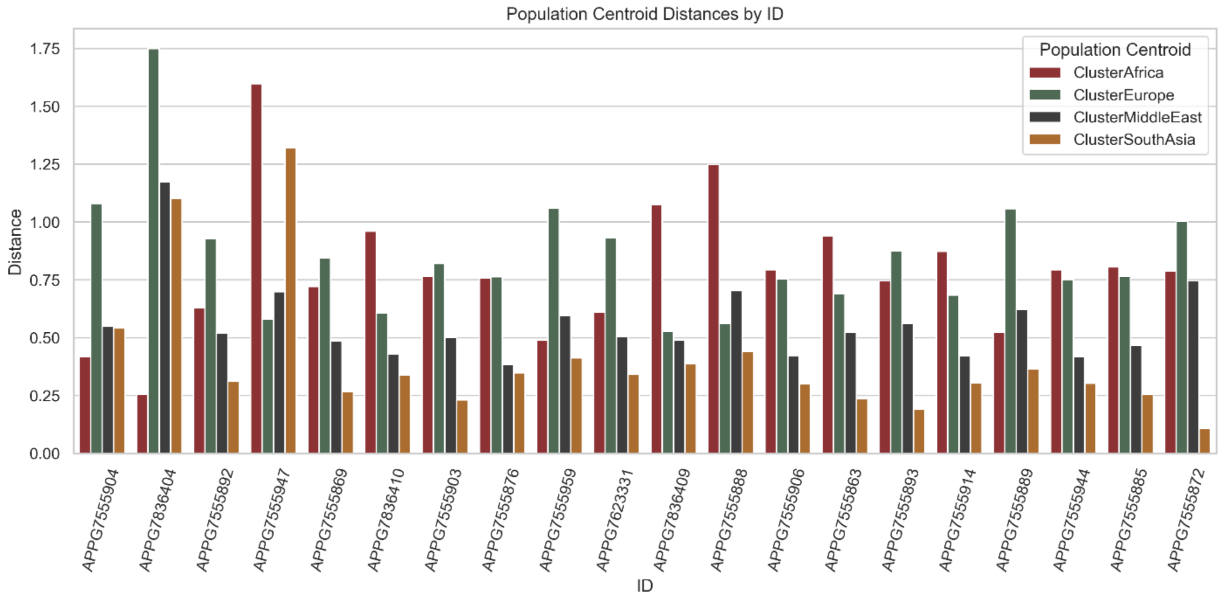


**Supplemental Figure 9** Distances to population centroids by ID for the 20 individuals where the Middle Eastern population centroid was not ranked the closest.

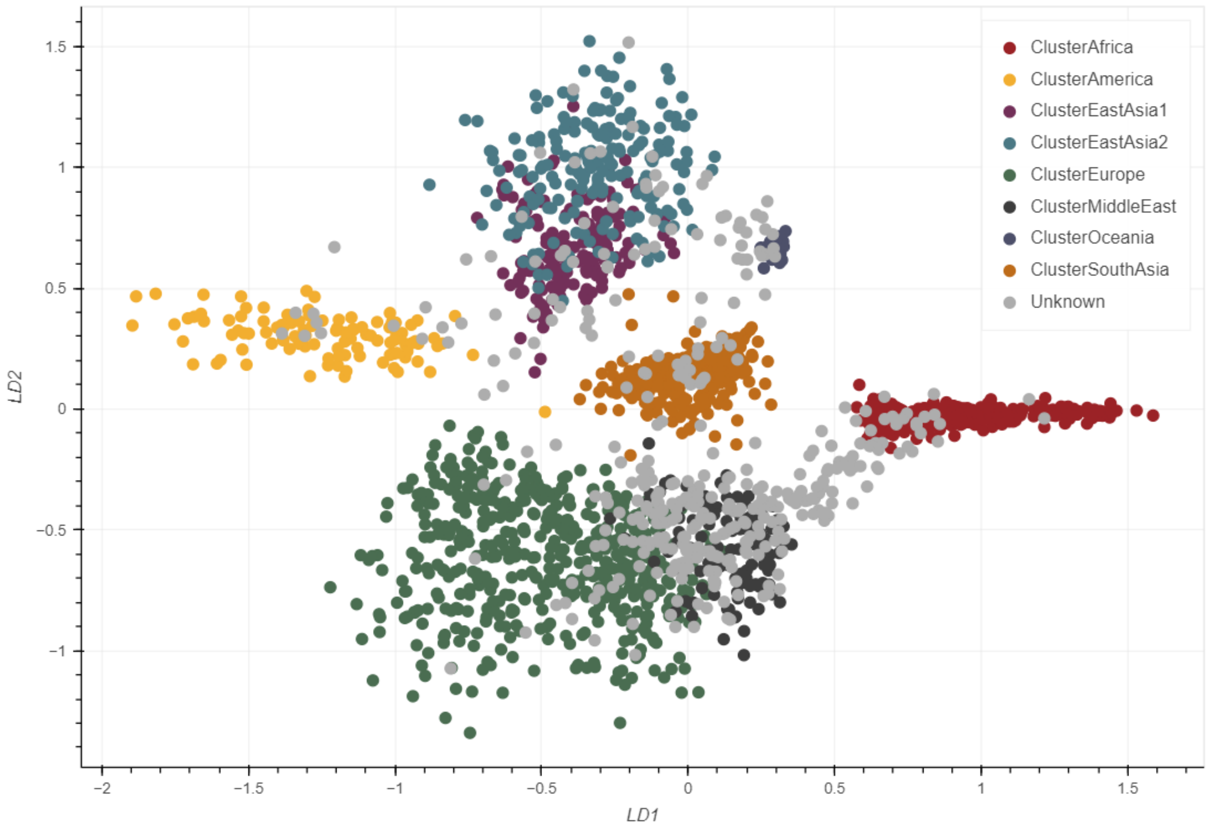


**Supplemental Figure 10** The selected samples (n=368) with over 90% Enhanced Tool coverage from Ruiz-Ramirez et al. (2023) labeled as “Unknown” when uploaded to our webtool that was developed as an adaptation of popVAE (Battey et al. 2021).
